# Evolution of constitutive bacteriocin production and release

**DOI:** 10.1101/2022.08.12.503745

**Authors:** Amrita Bhattacharya, Curtis M Lively

## Abstract

The production of the bacterial anti-competitor toxins, called bacteriocins, is widely described as spite, because the production of the toxins is costly and often requires cell death for release. Given these costs, it reasonable to think that bacteriocin production should be induced by the presence of unrelated competitor strains. Nonetheless, recent evidence in the insect-pathogenic bacterium *Xenorhabdus koppenhoeffri* showed that bacteriocin production occurs regardless of the presence of unrelated competitors. Could the constitutive production of bacteriocins be favored by natural selection? Here we use a mathematical model to examine this question, both within and between subpopulations. The model assumes that bacteriocin production is constitutive and costly, and that toxin release only occurs during natural cell death. We found that constitutive bacteriocin producers can outcompete non-producer sensitive strains within populations, and that it can spread in a metapopulation provided there is local competition for patches. Hence, the evolution of bacteriocin production does not require the detection of competitors or “suicidal” release of toxins.

## Introduction

The evolution and maintenance of spiteful behaviors that reduce the fitness of both the actor and the recipient pose a long-standing evolutionary puzzle (Hamilton, 1964, 1970). How can a trait that reduces the fitness of the actor be favored by natural selection? Nonetheless, bacteriocins are produced by almost all known lineages of bacteria (Klaenhammer, 1988; Riley & Chavan, 2006). Bacteriocins are proteinaceous toxins with diverse mechanisms of action, that can kill closely related local competitors without killing clone-mates (Reeves, 1965; Riley & Chavan, 2006). Bacteriocin production is considered spiteful because these toxins kill their targets and impose fitness costs of production, thus reducing the fitness of both recipients and actors (Gardner et al., 2004; West et al., 2007). In addition to the costs of DNA replication, maintenance, and protein synthesis, the release of these large, proteinaceous toxins often requires cell lysis, particularly in gram-negative cells that lack a cell wall (Riley & Wertz, 2002; Riley & Chavan, 2006; Wloch-Salamon et al., 2008).

Theory predicts that the competitive environment is key for the maintenance of costly bacteriocin production (Chao & Levin, 1981; Gardner et al., 2004). Bacteriocin-production is predicted to be most favored when competitors occur at comparable frequencies to the producing lineage (Gardner et al., 2004; Inglis et al., 2009). Further, a recent meta-analysis revealed that bacteriocin production is more likely to be associated with competition-related stressors such as starvation or cellular damage, than abiotic stressors such as heat or osmotic stress (Cornforth & Foster, 2013). This led to the proposition of the ‘competition sensing hypothesis,’ which posits that bacterial populations have evolved the ability to sense and respond to competition in their environment by upregulating bacteriocin production (Cornforth & Foster, 2013). While this may be generally true, the gram negative bacteria, *Xenorhabdus koppenhoeffri*, was found in vitro to produce and release the same levels of bacteriocins in both the presence and absence of non-self competitors (Bhattacharya et al., 2018). Why do *X koppenhoeffri* produce and release bacteriocins in the absence of competitors?

Here we address this question by building a mathematical model to ask whether the constitutive production of bacteriocins can be selected in natural bacterial populations. The model examines competition between bacteriocin-producer and bacteriocin-sensitive cells when bacteriocin production is constitutive and costly. We assume that toxins are only released upon natural cell death. Our results demonstrate that constitutive bacteriocin-producers can outcompete faster growing sensitive cells within patches, and that they increase in frequency in structured populations, if there is some degree of local competition.

## Model

We develop a model that tracks competition between two pathogenic bacterial strains: a bacteriocin producing strain and a sensitive strain that is killed by the producer’s bacteriocins. The strains compete for a finite amount of a non-renewable resource within a shared host. We assume that the production of bacteriocins is constitutive and costly. In other words, we assume that bacteriocin production is not a plastic response to the presence of competitors and that it imposes a fitness cost by reducing the birth rate of producers. Finally, we assumed that the release of bacteriocins only occurs during natural cell death, independent of whether competing cells are present. Importantly, the producer death rate is fixed and independent of competitor densities enabling the test of inducibility being necessary for the maintenance of costly bacteriocin production. First, we ask whether constitutive bacteriocin production could be favored by selection within hosts. We then examine whether constitutive bacteriocin production could increase when rare in a metapopulation in which there is between-host competition.

### Within-Host Competition

Let *B*_*t*_ be the “birth” number per sensitive cell at time *t*. We assumed *B*_*t*_ was both density dependent and resource dependent, as follows:

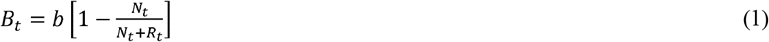

where *b* is the intrinsic number of new cells produced per bacterial cell per time step, *N*_*t*_ is the total number cells (producer + sensitive) in the population at time *t*, and *R*_*t*_ is the number of resource units remaining in the population at time *t*. The variable *b* was set to 1 in our simulations. We also assumed that each cell division requires one unit of the resources available at time *t*. Similarly, the death rate, *D*_*t*_, was also density-dependent and calculated as:

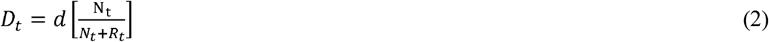

where *d* is the intrinsic death rate. The number of producer cells at time step, *t*+1, is then given by

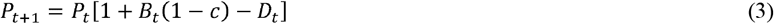

where *P*_*t*_ is the number of producer cells at time step *t*, and *c* is the birth-rate cost of bacteriocin production.

Toxin molecules were released only upon the natural death of producer cells. The total number of toxin molecules in the system at time *t*+1 was

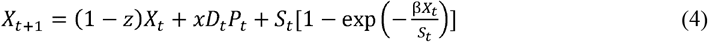

Where, *X*_*t*_ is the number of toxin molecules in the system at time *t*; *z* is the loss probability of toxin molecules; *x* is the number of toxin molecules released by each dying producer cell; and β is the contact probability between toxin molecules and sensitive cells, where *S*_*t*_ is the number of sensitive cells at time *t*. The first term on the right-hand side (RHS) of eq. (4) thus gives number of toxin molecules carried over from the previous time step. In the present study we set *z* equal to 0.05 to give a 5% decay rate. The second term on the RHS gives the number of new toxin molecules released into the system resulting from producer deaths. The third term on the RHS of (4) gives the number of sensitive cells that were killed by contact with toxin, where *x*_*t*_*/S*_*t*_ gives the mean number of toxin molecules per sensitive cell. Note that the exponential term 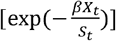 gives the probability of the zero class in a Poisson distribution; hence 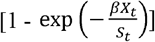 gives the probability that a sensitive cell would contact at least one toxin molecule, thus killing the sensitive cell. Thus, the third term on the RHS gives the number of toxin molecules removed from the system by killing sensitive cells, assuming each toxin-induced death removes a single toxin molecule from the system.

The number of sensitive cells at time step *t*+1 was calculated as:

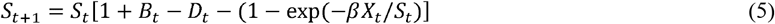

where *S*_*t*_ is the number of sensitive cells at time *t*. After every time step, the number of resource

units remaining in the system were calculated as:

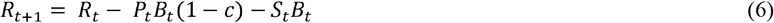

where *R*_*t*_ is the number of resource units at time t. The model thus assumes that the birth of every cell requires one resource unit.

The recursion equations above were iterated using R (R Core Team. 2019). The code is given in Supplementary Files 1 and 2. The simulation was initiated with 10 producer cells and 90 sensitive cells in a single host containing 10^8^ resource units. We ran the simulation for 100 time steps, which was more than enough for the resources to be depleted to zero. The intrinsic birth rate was set to 1 (*b* = 1), and the intrinsic death rate was set to 0.01 (*d* = 0.01). The cost of toxin production was set to 0.05 (*c* = 0.05). We ran the model for a range of parameters for β and *x* to explore the effects of the contact probability between toxin and sensitive cells, and the number of bacteriocin particles released following the death of producer cells.

### Between-Host Competition

For bacteriocin production to be favored by natural selection, producers not only have to “win” within hosts, but they also require an advantage under some level of between-host competition. Such an advantage likely depends on the degree of soft selection and migration. Here we used Frank’s “scale-of-competition” method to examine bacteriocin evolution in a metapopulation (Frank, 1998). The method uses the variable ‘*a*’ to represent the probability of competition with individuals derived from the same patch. The variable can also be seen as a combination of soft selection (s) and migration (m)(van Dyken, 2010), where

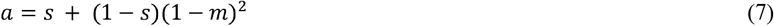

The model assumes a metapopulation comprising of a large number of hosts where each host is colonized by a fixed initial number of bacteria (*N* = 100). One host in the metapopulation is colonized by a mixture of 10 producer cells and 90 sensitive cells. This is called the “local” host. All remaining hosts are colonized by 100 sensitive cells.

The relative fitness of bacteriocin producers (*W*_prod_) was calculated as (Frank, 1998):

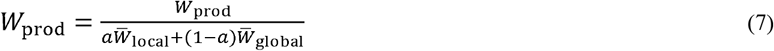

where, *W*_prod_ is the number of producer cells at time *t* (*P*_*t*_) divided by the number of colonizing producer cells (*W*_prod_ = *P*_*t*_*/P*_1_) where *P*_1_ =10; 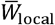 is the number of cells per colonizing cell in the local host (i.e., (*P*_*t*_ *+ S*_*t*_)/100) containing the producer cells; and 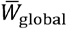 is the number of cells per colonizing cell in the non-focal hosts at dispersal, i.e., *S*_tot_/100, where *S*_tot_ is the total number of sensitive bacteria in non-local hosts at dispersal.

## Results

### Within-Host Competition

The model tracks the densities of bacteriocin-producer cells, bacteriocin-sensitive cells, and resource units within the host under varying contact probability between toxin molecules and sensitive cells, *β*, and varying levels of the number of bacteriocin particles released per dying producer cell, *x* (Figure 1). Each simulation begins with 10 producer cells and 90 sensitive cells within a shared host. When *x* is set to 0 (Fig 1 a-c) the densities of bacteriocin producers never exceed the densities of bacteriocin sensitive competitors, demonstrating that producers are unable to outcompete sensitive cells. However, bacteriocin-producing cells outcompete bacteriocin-sensitive competitors when *x* is non-zero (Fig 1 e-i). Producers outcompete sensitive cells faster at higher levels of contact probability, β, and with higher numbers of bacteriocin particles released per dying producer cell, *x* (Fig 1).

**Figure 1:**
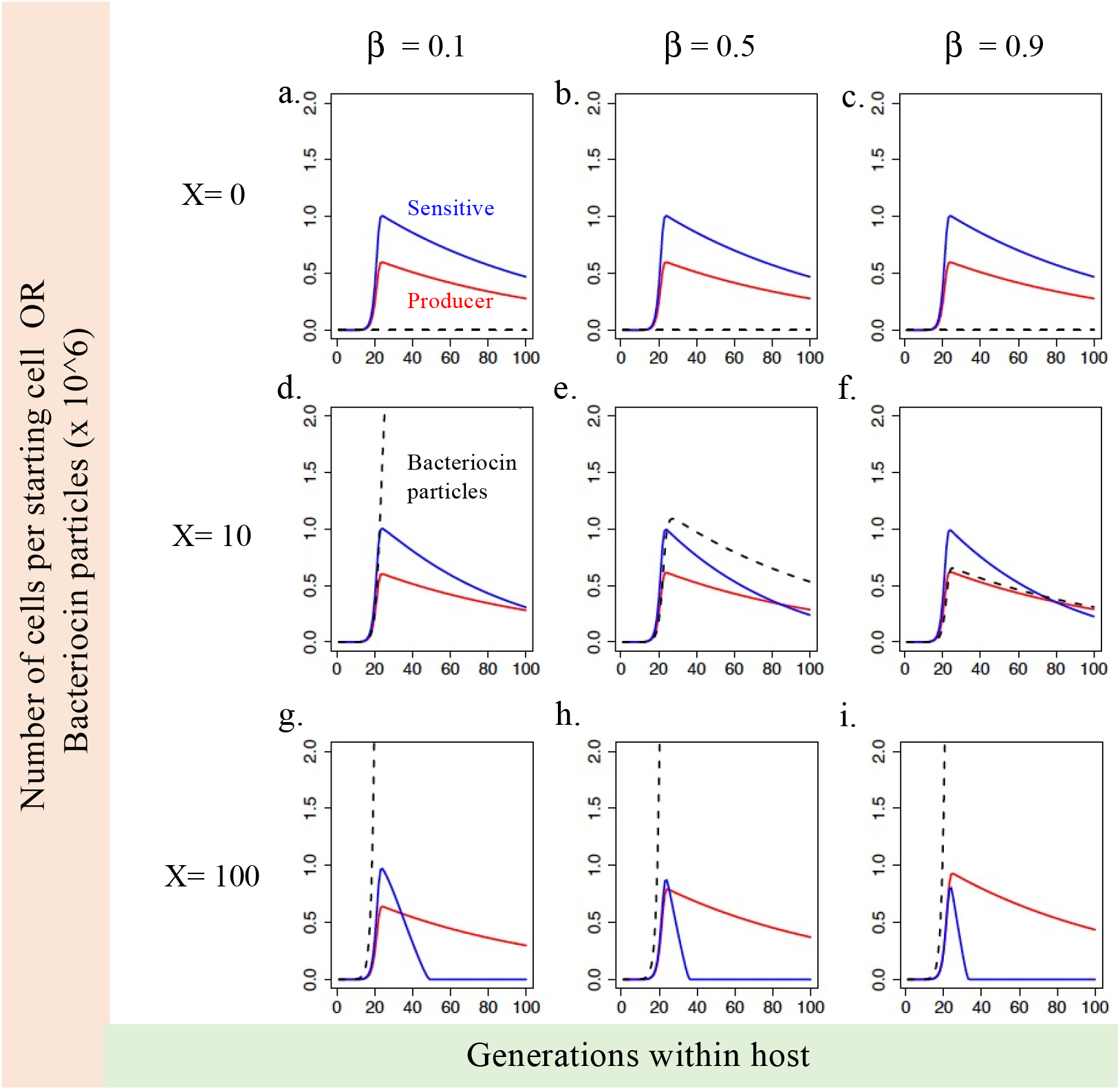
Within-host competitive dynamics of bacteriocin producing cells (red) and bacteriocin sensitive competitors (blue). Red and blue lines represent the number of producer or sensitive bacterial cells, respectively, relative to the initial number of cells of each type. Variable *x* represents the number of bacteriocin particles released by every dying producer cell. Variable β represents the contact probability between bacteriocin particles and sensitive cells. Black dashed lines represent the total number of unused bacteriocin particles in the environment. Panels a-c represent competitive dynamics when no bacteriocin is not effective at killing sensitive cells. Here sensitive cells outcompete producer cells every time and the slower producer growth represents the growth cost of bacteriocin production incurred by producer cells. Producer cells can outcompete sensitive cells when *x* > 0 and do so faster with increasing contact probability β between bacteriocins and sensitive cells (panels e – i).

### Between-Host Competition

Relative fitness of bacteriocin producers is examined across a metapopulation of hosts under initial conditions of *β* = 0.1, varying levels of *x*, and three difference scales of competition (Fig 2). Relative fitness measures greater than 1 indicate invasion of the metapopulation by producers despite being rare initially. Our results demonstrate that the relative fitness of bacteriocin producers can exceed 1 under many conditions (Fig 2 d-i) despite conservative assumptions about bacteriocin potency (β = 0.1; Fig 2). Invasion occurs faster and is favored more strongly at more local scales of competition i.e., higher values of ‘*a*’, and with higher numbers of bacteriocin particles released per dying producer cell, *x*.

**Figure 2:**
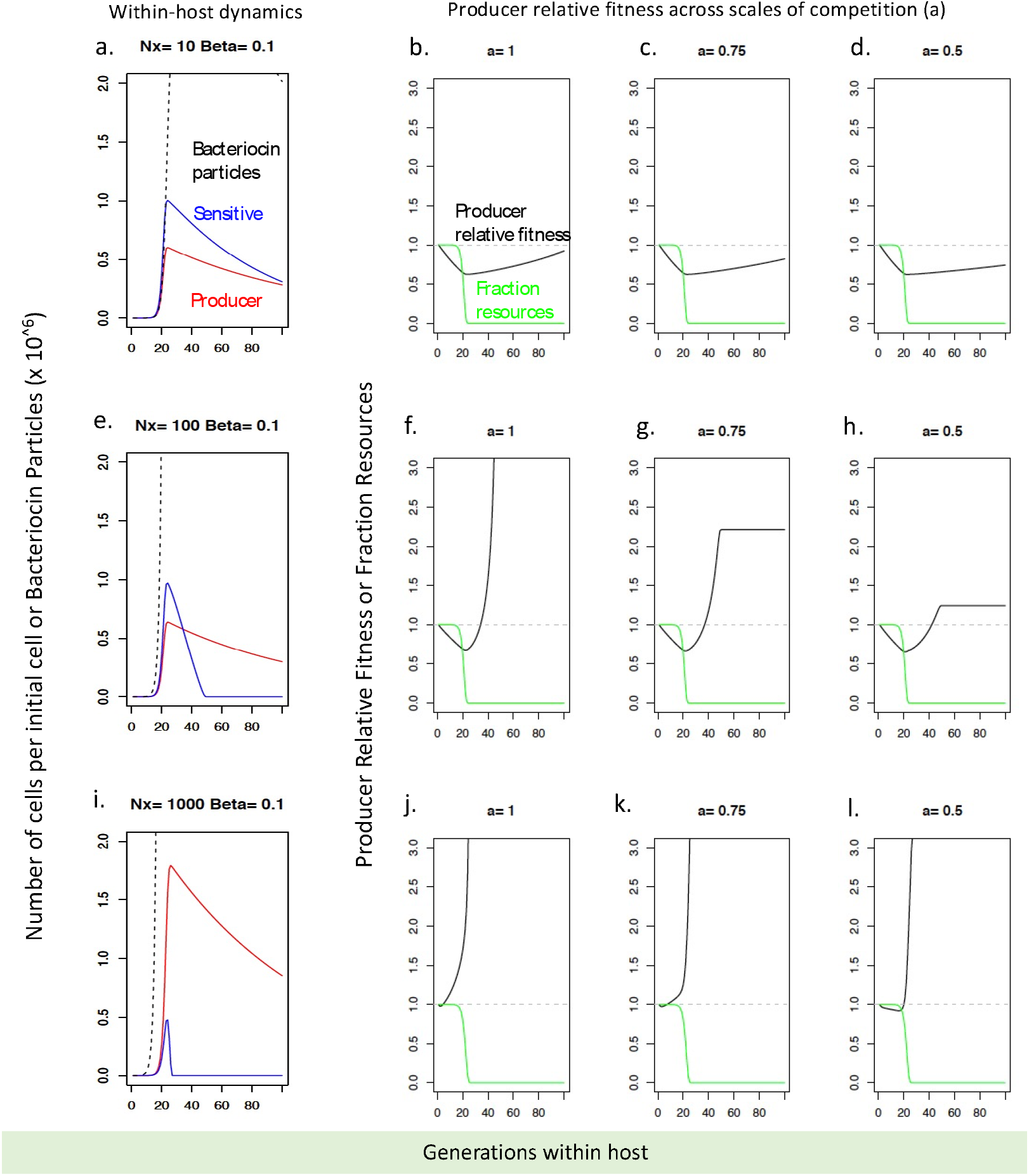
Within-host dynamics and relative fitness of bacteriocin producer cells across the metapopulation of hosts. Fig 2 a, e and i represent the within-host growth of bacteriocin producers and sensitive competitors for varying number of bacteriocin particles released per dying producer cell (X). Panels 2 b-d, f-h, j-l show the relative fitness of producer cells across the metapopulation of hosts for varying values of scale of competition, ‘a’ (Frank, 1994). Scale of competition, a can range from completely local (a = 1) to completely global (a =0).

## Discussion

Bacteriocin production is an example of spiteful behavior, meaning that both the actor and the recipient pay fitness costs (Riley & Chavan, 2006). Bacteriocin production is frequently considered a ‘suicide mission’ where producer cells sacrifice themselves to deploy these anti-competitor toxins (West et al., 2007; Nedelcu et al., 2010). Inducible mechanisms that allow bacteriocin production to be increased by the competitive environment are predicted to be favored by selection (Gardner et al., 2004; Cornforth & Foster, 2013). Yet, contrary to expectations, bacteriocin production in *X koppenhoeffri* was found to occur even in the absence of non-self competitor cells (Bhattacharya et al., 2018). Our goal here was to understand why. We built a mathematical model to examine if and how, despite considerable costs of production and release, bacteriocin-producing lineages like *X. koppenhoeffri* persist.

Our results demonstrate that a bacteriocin-producing lineage can outcompete a faster growing sensitive lineage, if bacteriocins are released only upon natural cell death (Fig 1 and 2). The time taken for the producing lineage to outcompete the sensitive lineage decreases with increasing transmission efficiency, β, and higher numbers of bacteriocin particles released per dying producer cell, *x* (Fig 1). Further, the results show that bacteriocin producers can invade a metapopulation of sensitive cells, provided some degree of local competition (Figure 2). Consistent with previous work on the scale of competition in general (Durrett & Levin, 1997; Gardner & West, 2004; Gardner et al., 2004), our results show that increasing local competition favors the invasion of bacteriocin production (Figure 2b). These results demonstrate that, despite cell deaths being necessary for bacteriocin release, inducible or higher-than-natural rates of cell death for ‘suicidal’ bacteriocin release are not required for bacteriocin-producing lineages to persist and increase in frequency when rare. Plasticity, however, could increase the advantage of bacteriocin production under some conditions (Bhattacharya, 2019; Granato & Foster, 2020; Niehus et al., 2021).

The assumptions about costs of bacteriocin production and bacteriocin potency (number of particles released per dying cell) in our models are probably conservative. The cost of bacteriocin production is set at 5% reduction in fitness (c = 0.05), which imposes a considerable growth disadvantage on the producer strain relative to the sensitive competitor. It has been previously argued that the fitness costs of bacteriocin production may be much less steep than commonly assumed (Dykes & Hastings, 1997). Empirical estimates in *E coli* that produce the most well-studied group of bacteriocins called colicins, suggest that every lysing producer cell is capable of releasing bacteriocin particles on the order of thousands of particles per cell (Gordon & Riley, 1999). Our model examines a wide range of this parameter ranging from 0-1000 particles of bacteriocin per dying producer cell, remaining highly conservative. Thus, our results demonstrate that the competitive benefits of bacteriocin production may be realized without invoking inducible mechanisms or ‘suicide’ despite high costs and low potencies of toxins produced.

It is important to note that these results do not preclude the possibility of plastic bacteriocin production or release. Indeed, recent work in *E coli* (Mavridou et al., 2018; Granato & Foster, 2020) provide evidence for increased toxin production in response to attack from a neighboring competitor. However, bacteriocin production in *E coli* has also been reported to occur in the absence of competitors, typically at late growth stages (Eraso et al., 1996). Additionally, recent work using models to compare various mechanisms of bacteriocin production and release found that toxin sensitivity to be the preferred mode of bacteriocin production and release over unregulated mechanisms (Niehus et al., 2021). Our goal here is not to claim that bacteriocin production cannot be inducible, rather to demonstrate that non-inducible mechanisms of bacteriocin release can maintain bacteriocin production. We aim to caution against the generalization of bacteriocin production as a ‘suicide’ mission and to encourage additional direct investigation of the inducibility of this spiteful, yet ubiquitous bacterial trait.

To summarize, the model described in this study demonstrates, using conservative assumptions, that inducible mechanisms of bacteriocin expression aren’t necessary to explain the maintenance of bacteriocin production in nature. Further, these results challenge the common assumption that bacteriocin production and release is a ‘suicide mission.’ Our models provide an alternative mechanism to explain the maintenance of spiteful bacteriocin production through canalized, constitutive expression instead of inducible production. We do not claim that canalized spite is the only, or even the most common, mechanism of bacteriocin production. However, we do offer a previously underexamined mechanism to explain the early evolution of bacteriocin production, and how spiteful bacteriocin production may be maintained in some species.

## Acknowledgements

We would like to thank the Center for Integrative Studies in Animal Behavior at Indiana University for fellowship funding to AB when this work was conducted. We also thank Mike Wade and Farrah Bashey for helpful comments on a previous draft

---

**Supplementary Material 1:**
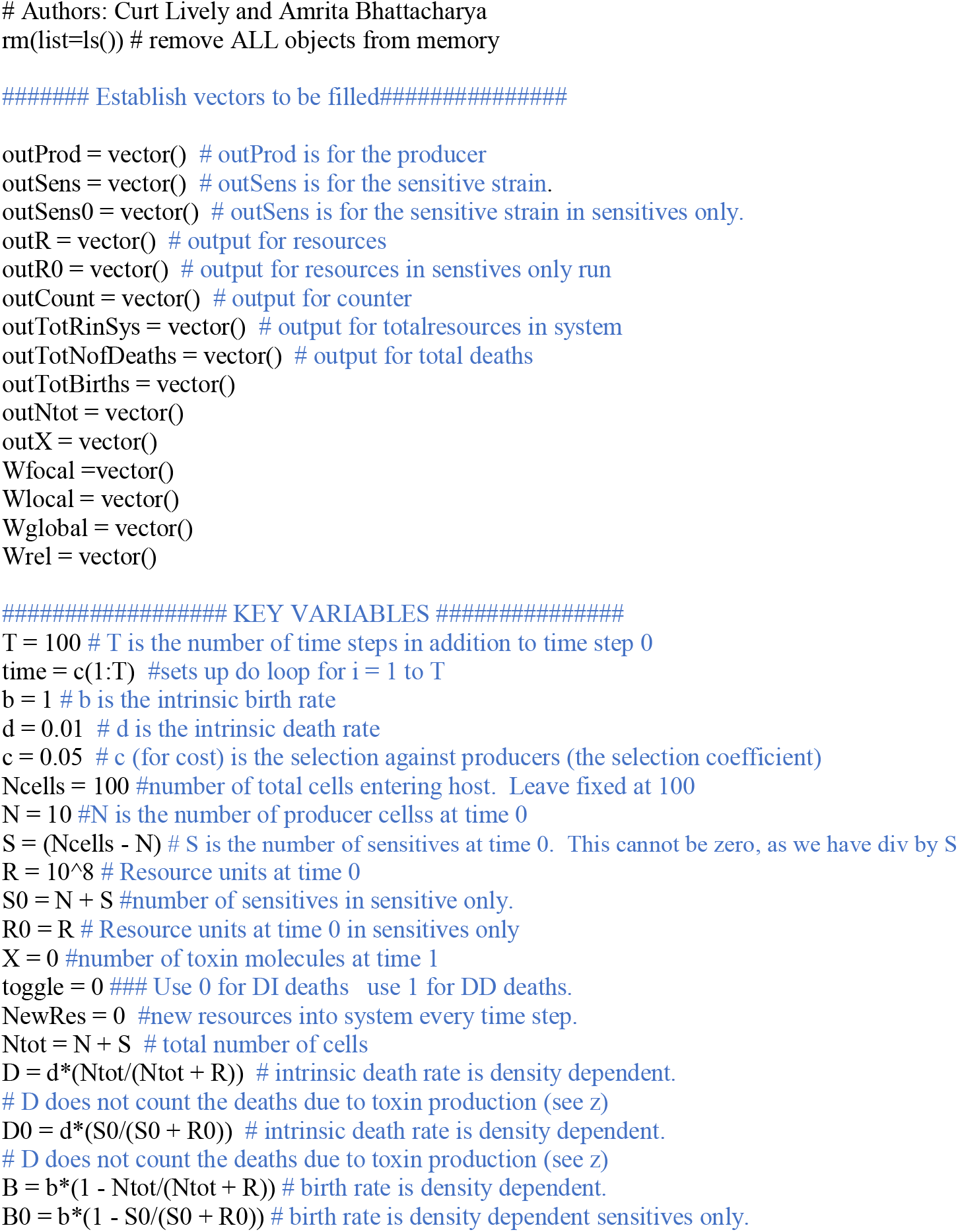
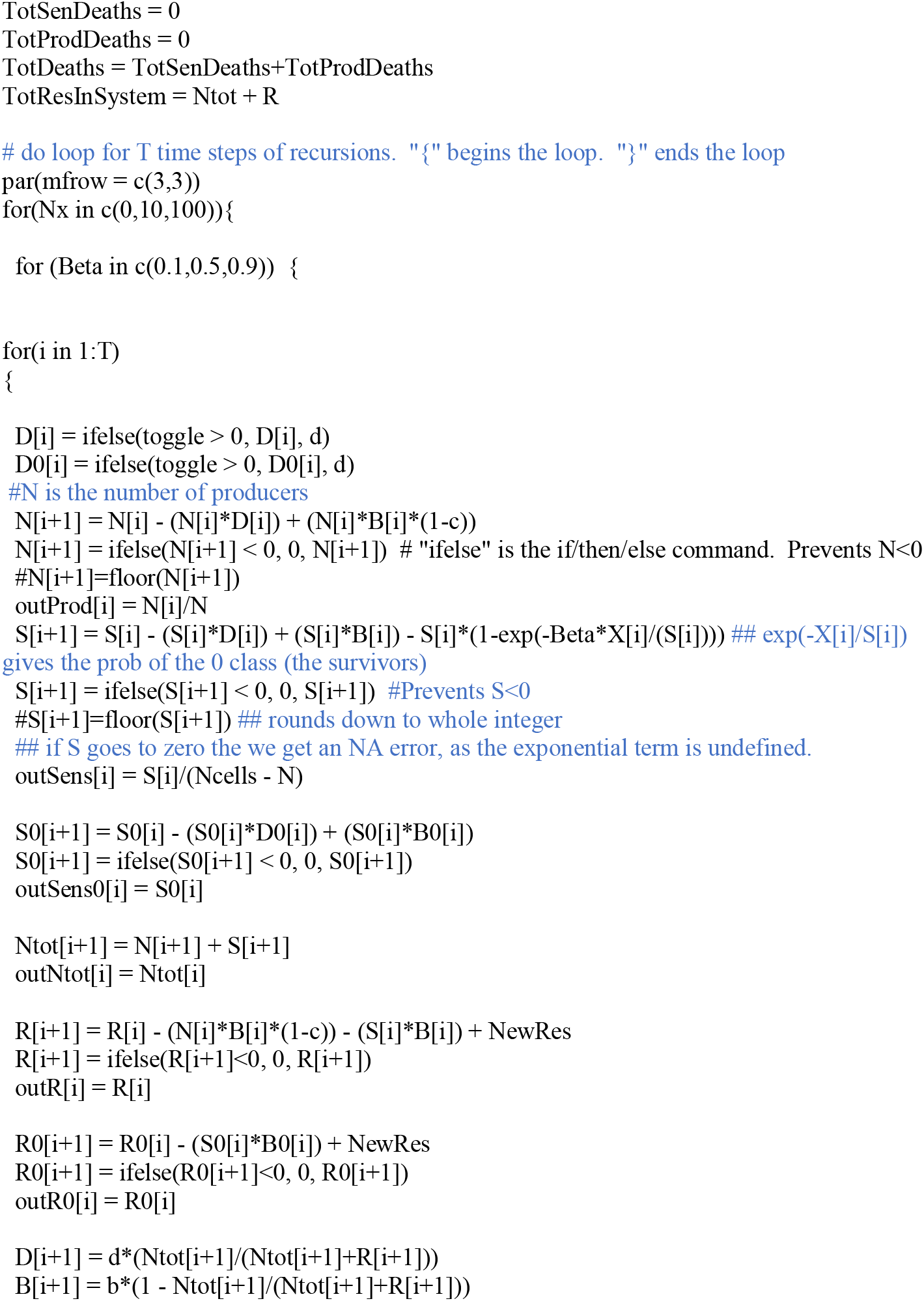
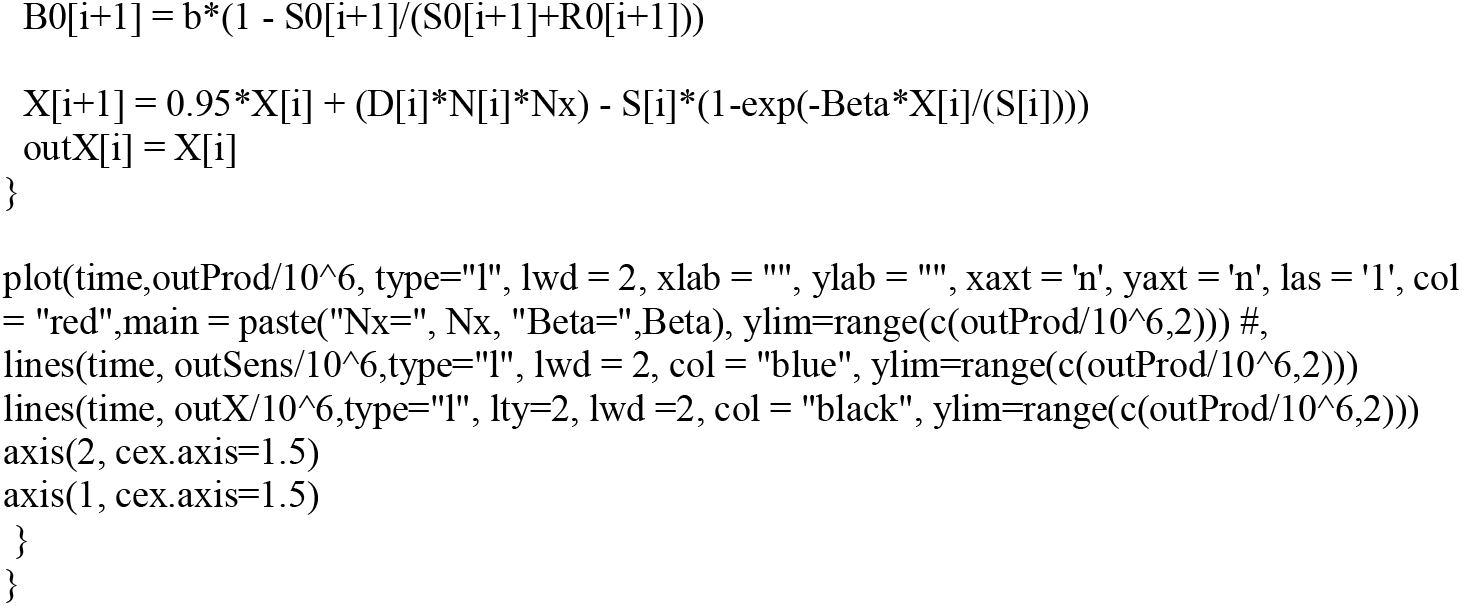
R code used to simulate within-host dynamics and produce figure 1.

**Supplementary Material 2.**
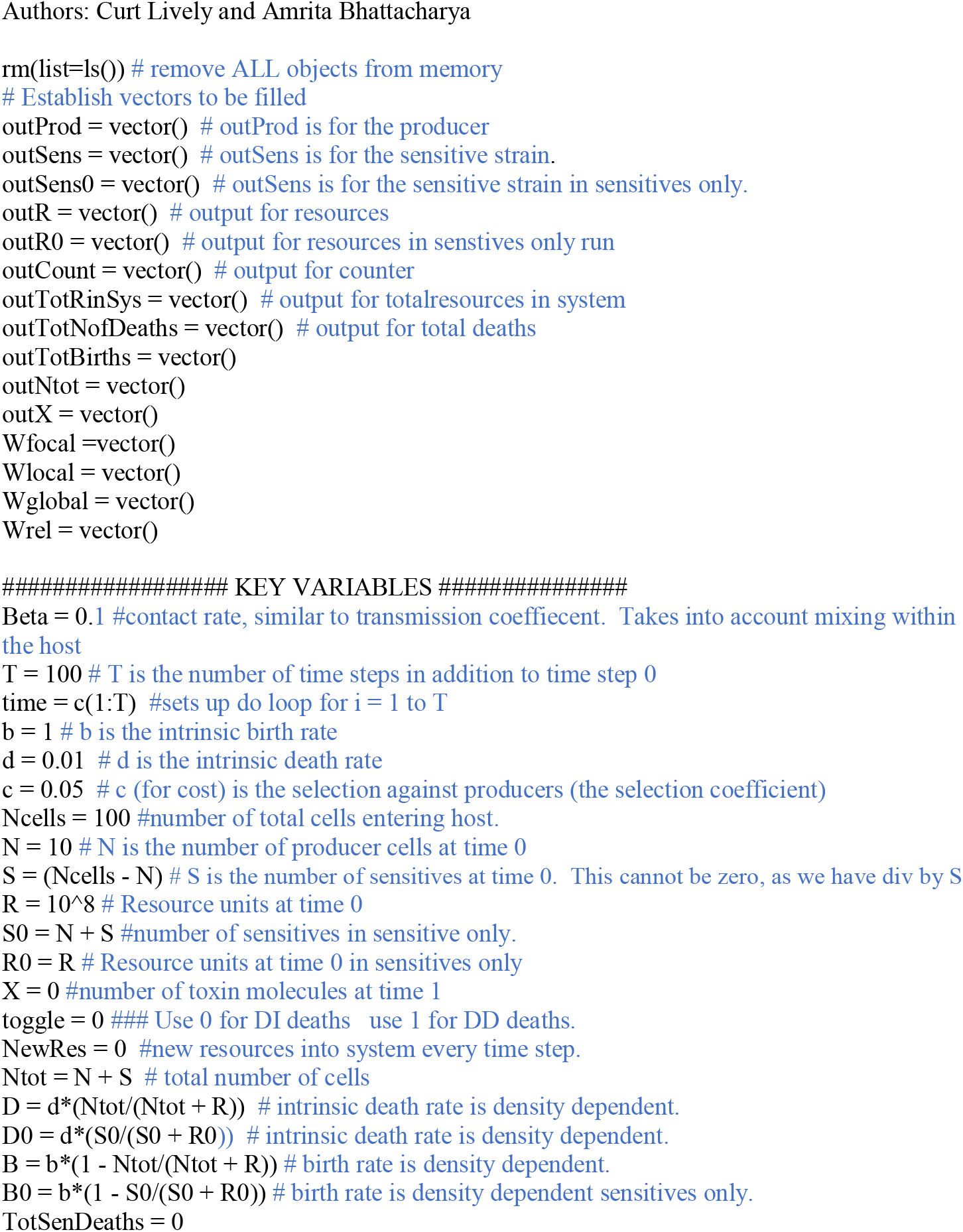
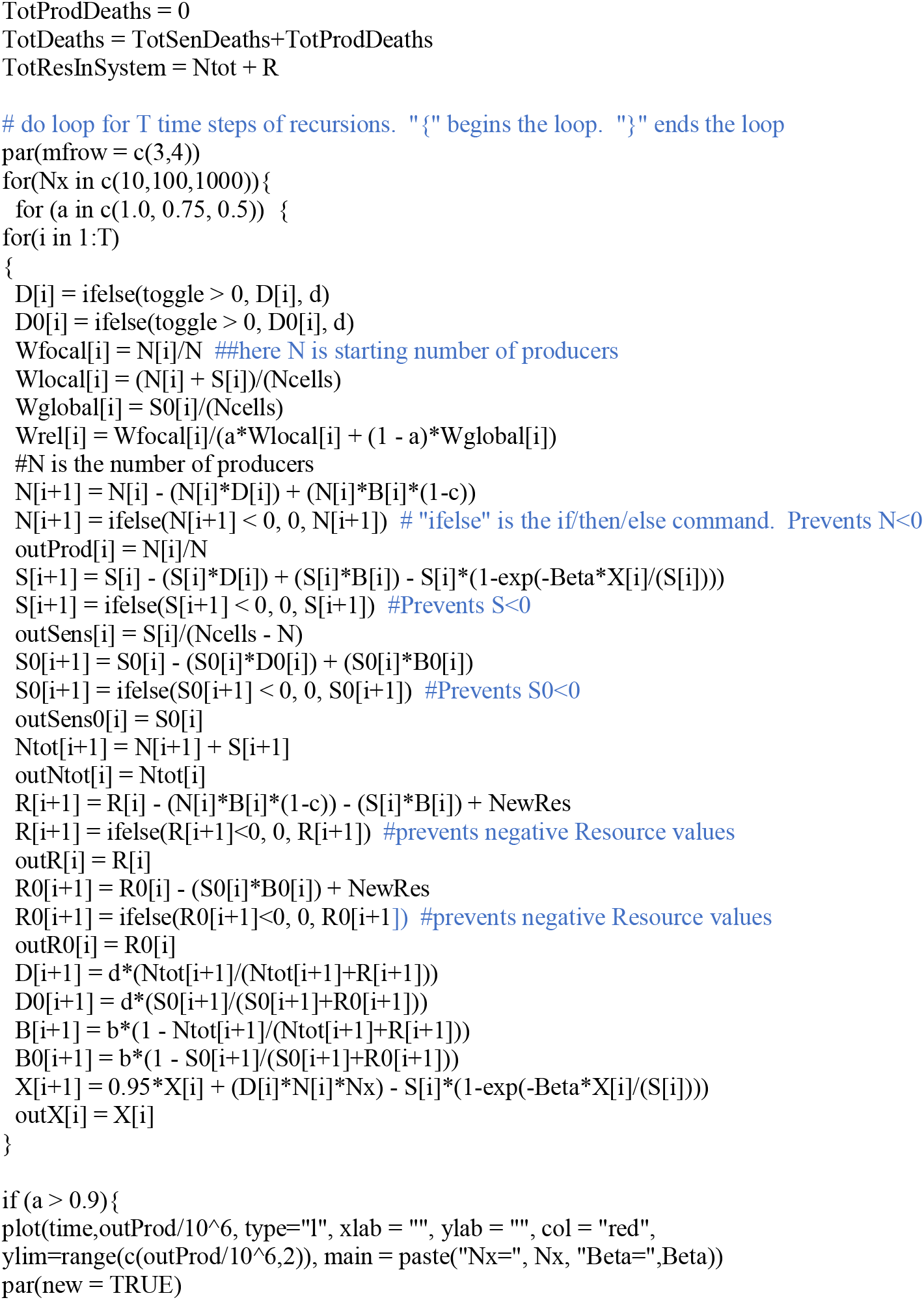
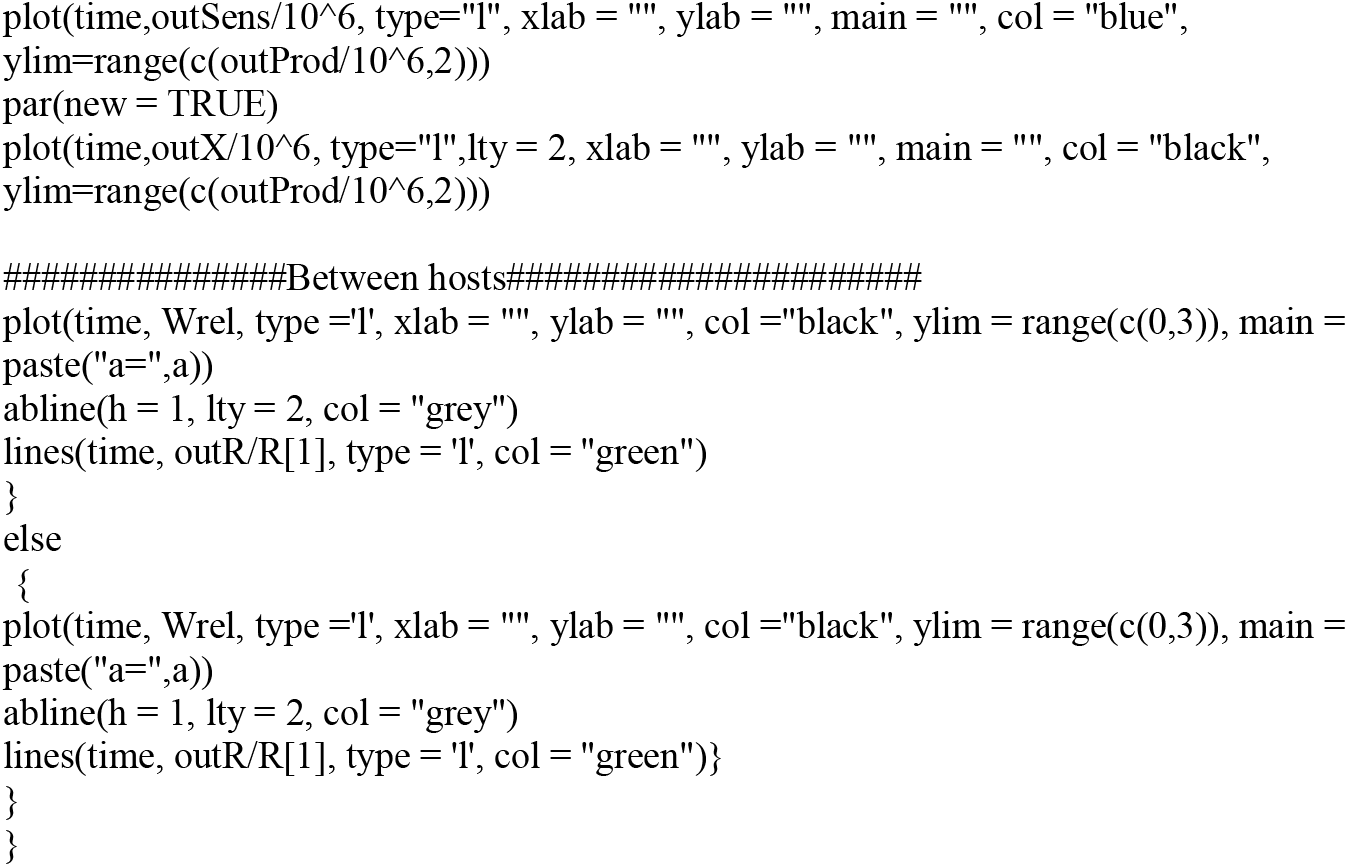
R code used to simulate between-host competition dynamics and produce figure 2.

